# Transcriptomic Profile of Lin^-^Sca1^+^c-kit (LSK) cells in db/db mice with long-standing diabetes

**DOI:** 10.1101/2024.01.22.576754

**Authors:** Neha Mahajan, Qianyi Luo, Surabhi Abhyankar, Ashay D. Bhatwadekar

## Abstract

The Lin^-^Sca1^+^c-Kit^+^ (LSK) fraction comprises multipotent hematopoietic stem cells (HSCs), vital to tissue homeostasis and vascular repair. While HSC homeostasis is impaired in diabetes, it is not known how chronic (>6 months) type 2 diabetes (T2D) alters the HSC transcriptome. Herein, we assessed the transcriptomic signature of HSCs in db/db mice employing mRNA and miRNA sequencing. We uncovered 2076 mRNAs and 35 miRNAs differentially expressed in db/db mice, including two novel miRNAs previously unreported in T2D. Further analysis of these transcripts showed a molecular shift with an increase in the pro-inflammatory cytokines and a decrease in anti-inflammatory cytokine expression. Also, pathway mapping unveiled inflammation and angiogenesis as one of the top pathways. These effects were reflected in bone marrow mobilopathy, retinal microglial inflammation, and neurovascular deficits in db/db mice. In conclusion, our study highlights that chronic diabetes alters HSCs’ at the transcriptomic level, thus potentially contributing to overall homeostasis and neurovascular deficits of diabetes, such as diabetic retinopathy.

**Highlights:**

- Bone marrow mobilopathy with long-standing diabetes
- Switch in LSK transcriptomic profile towards inflammation and angiogenesis
- Discovered 35 miRNAs, including two novel miRNAs, miR-3968 and miR-1971
- LSK dysfunction reflected in inflammation and neurovascular deficits of the retina

## Introduction

Diabetes is a chronic debilitating metabolic disease affecting more than 537 million people worldwide, and this number is predicted to rise to 783.2 million by 2045 (Sun et al., 2022) (IDF, 2021). Hematopoietic stem cells (HSCs) are bone-marrow-derived stem cells that can differentiate into myeloid, lymphoid cells, platelets, and red blood cells (Hanoun and Frenette, 2013). These bone marrow-derived HSCs are adversely affected by chronic diabetes. With the primary role of tissue maintenance, homeostasis, and vascular repair, these HSCs in diabetes lose their ability to be reparative, instead, it has been reported in multiple studies that mobilopathy, stem cell rarefaction, neuropathy, micro-angiopathy, and inflammation become more prevalent in HSCs with diabetes (Fadini and Albiero, 2022; Vinci et al., 2020). Diabetes also manifests many micro- and macro-vascular complications, such as retinopathy, neuropathy, cardiomyopathy, and nephropathy, posing a high risk for multi-organ damage (Paul et al., 2020). Consistently, it has been shown that low levels of HSCs in chronic diabetes accelerate the progression of diabetes-induced microvascular complications (Albiero et al., 2020; Fadini et al., 2017).

The transcriptomics studies have strengthened our knowledge of genetic factors associated with the development and associated outcomes of diabetes. Importantly, different groups have utilized the insulin-targeted organs, such as the pancreas, liver, skeletal muscles, adipose tissues, etc., for gene expression studies reflecting the dominance of inflammatory immune responses in the pathogenesis of diabetes (Corbi et al., 2020; Lin et al., 2020; Tonyan et al., 2022); however, LSK transcriptome under long-standing diabetes has not been studied well. In addition to mRNA targets, non-coding RNAs like micro RNAs (miRNA) have been widely explored (He et al., 2021). Recent evidence has shown the potential of tissue-specific and circulating miRNAs for their capability to be used as a biomarker in diabetes and associated vascular complications (Liang et al., 2018; Smit-McBride and Morse, 2021). Therefore, studying miRNA and mRNA transcriptomic could provide critical information about LSK cell health under conditions of long-standing diabetes.

It is noteworthy that HSC dysfunction is a known outcome of diabetes (Fadini and Albiero, 2022; Vinci et al., 2020); however, the transcriptome signature of these HSCs in diabetes has not been investigated. Additionally, it is still unknown if and how the transcriptomic alterations of HSCs with diabetes can contribute to vascular dysfunctions. In the present study, we isolated a highly purified fraction of HSCs; Lin^-^Sca1^+^c-Kit^+^ (LSK) cells from the bone marrow of db/db mice (an animal model of type 2 diabetes) with six months of diabetes followed by miRNA and mRNA sequencing. Simultaneously, we used the retina as a tissue to study the manifestations of chronic diabetes and the impact of HSC health because diabetic retinopathy is the most common complication of diabetes, and the intricate nature of retinal neovasculature provides a unique window to study the effect of diabetes to understand the pathology of higher centers such as the brain. Our studies unravel previously unknown miRNAs and mRNA transcripts changed with long-term diabetes, which could have contributed to HSC mobilopathy and neurovascular deficits of the retina.

## Results

### The bone-marrow-derived HSCs and diabetes

Diabetes is known to dysregulate the bone marrow (Vinci et al., 2020), but very little is known about the transcriptional dysregulation of HSCs in long-term diabetes. Therefore, we evaluated the pure population of HSCs using miRNA and mRNA sequencing (**Figure 1A**). The bone marrow was first subjected to flow cytometric sorting to isolate the lineage^neg^Sca1-ckit^+^/ LSK cells. Assessment of the pure LSK cell population showed an increase in db/db mice (**Figure 1B**), indicating bone marrow mobilopathy with chronic diabetes. The pure population of LSK cells was then used for the transcriptome analysis. Out of 13708 genes with non-zero total read count; 2076 genes were significantly expressed in HSCs of db/db mice (p <0.05) **(Figure S1A)**. The volcano plot represents the total genes identified (**Figure S1B**), whereas the heatmap in **Figure 1C** represents significantly changed genes. Of these, 1036 were upregulated, whereas 944 were downregulated in the db/db mice (**Figure S1A**).

**Figure 1.**
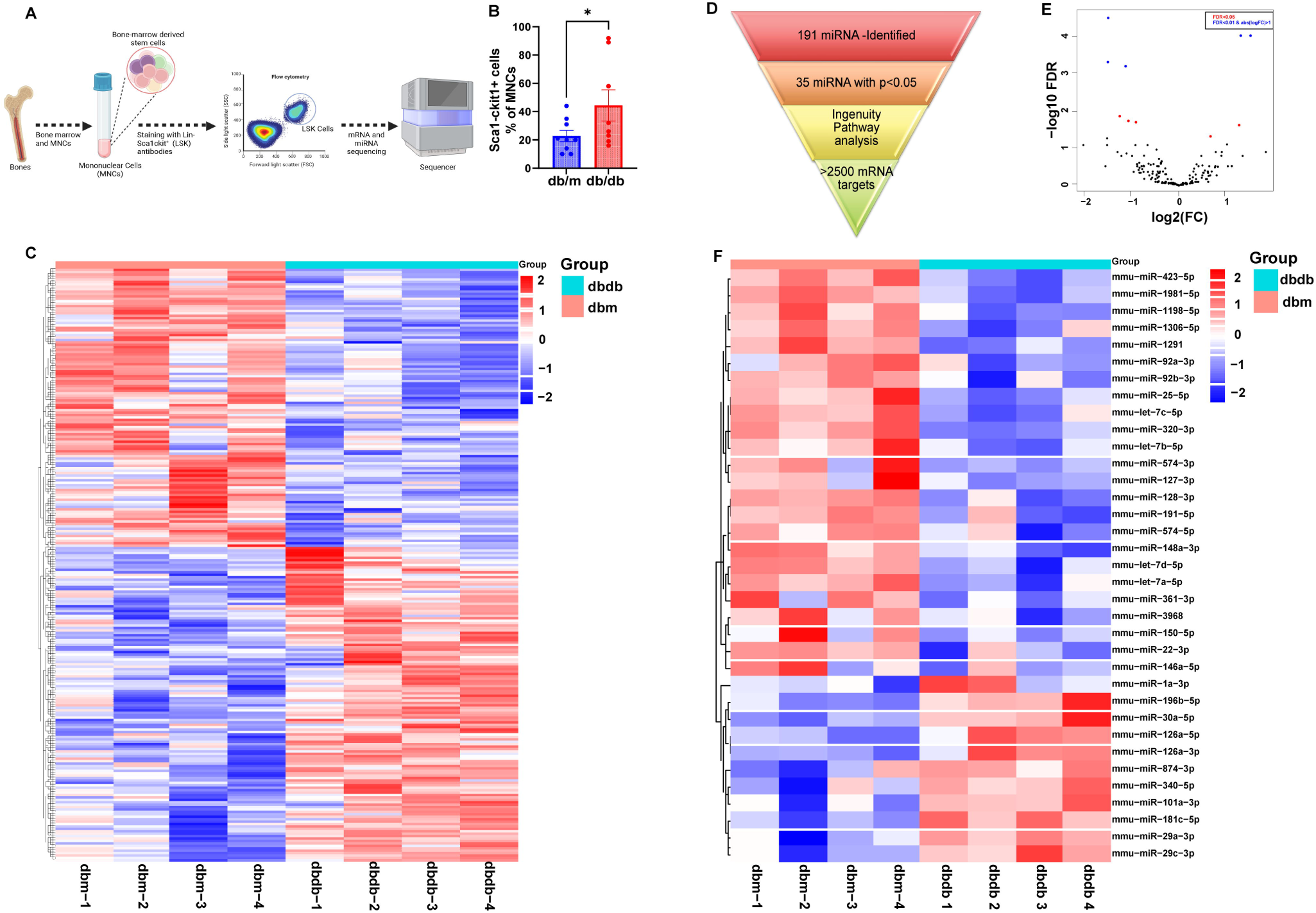
Transcriptome analysis of diabetic HSCs. **A)** Schematic presentation of the steps followed for isolating, sorting, and sequencing HSCs from the bone marrow. **B)** The sorted HSCs population was found to be higher in the db/db mice. **C)** Heatmap depicting the significantly changed mRNAs between db/m and db/db mice. **D)** The representative procedure for the miRNA data analysis. **E)** The volcano plot for the total miRNAs found in the sequencing data. The red and blue dots represent the significantly changed miRNAs. **F)** The top 35 miRNAs are presented as a heatmap. The data in Figure **(B)** is presented as mean ± SEM, analyzed using student unpaired t-test; N=9. The heatmaps in Figures **(C)** and **(F)** were plotted using a freely available online tool SRplot (https://www.bioinformatics.com.cn/en).

miRNA sequencing found a total of 191 miRNA, out of which 35 miRNAs were significantly changed in db/db mice (p<0.05) (**Figure 1D**). **Figure 1E** represents the volcano plot for a total of 191 miRNAs observed from the sequencing, whereas the top 35 miRNAs were presented in a heatmap in **Figure 1F**. The differentially expressed miRNAs (35) were then subjected to Ingenuity pathway analysis (IPA) for a deeper understanding of their involvement in various molecular pathways. The core analysis from IPA showed the top 10 upregulated and downregulated miRNAs in db/db mice (**Table S1**). Micro RNA target filter analysis *via* IPA software showed that out of 35 significantly changed miRNAs, 28 miRNAs had 1974 mRNA targets. Further, these mRNA targets were traced back to the mRNAs identified in the LSK cells of the present study, which revealed that 446 mRNAs were also present in our mRNA sequencing data, on the other hand, 1528 genes or mRNA targets were unique or were absent in our sequencing data (**Figure S1C**). These miRNAs had several downstream targets, some unique to the individual miRNA, and some overlapping between multiple miRNAs (**Figure S1D**).

### Transcriptome analysis of bone-marrow-derived LSK cells in chronic diabetes

Gene ontology using the online available software Panther was performed to investigate the molecular pathways associated with the differential mRNAs in the LSK cells. It was found that most of the genes were associated with inflammation mediated by chemokine and cytokine signalling pathways **(Figure 2A)**. Interestingly, angiogenesis was also one of the top pathways having a significantly higher number of genes in the transcriptome of db/db mice’s LSK cells **(Figure 2A)**. A significant increase in the angiogenic genes, such as *Vegfc, Pdgfd, Angpt1, Tgfb1i1, Pla2g4c, and Hif3a* **(Figure 2B)** was found in the diabetic LSK cells. We also found an increase in the expression of *Cxcl12, Cxcl5, and Csf1* **(Figure 2C)**. No significant change was observed in the expression level of *Cxcr4* in LSK cells **(Figure 2C)**. To study inflammatory changes, we investigated our mRNA data for major inflammation-related genes, including Toll-like receptors (TLRs), interleukins (ILs), and tumour necrosis factor (TNF). The anti-inflammatory cytokine *Il10* was significantly downregulated in the HSCs of db/db mice (**Figure 2D)**. In contrast, a significantly higher expression of *Il4, Tlr4,* and *Tnf11*α (**Figure 2E)** was found in the HSCs of db/db mice.

**Figure 2.**
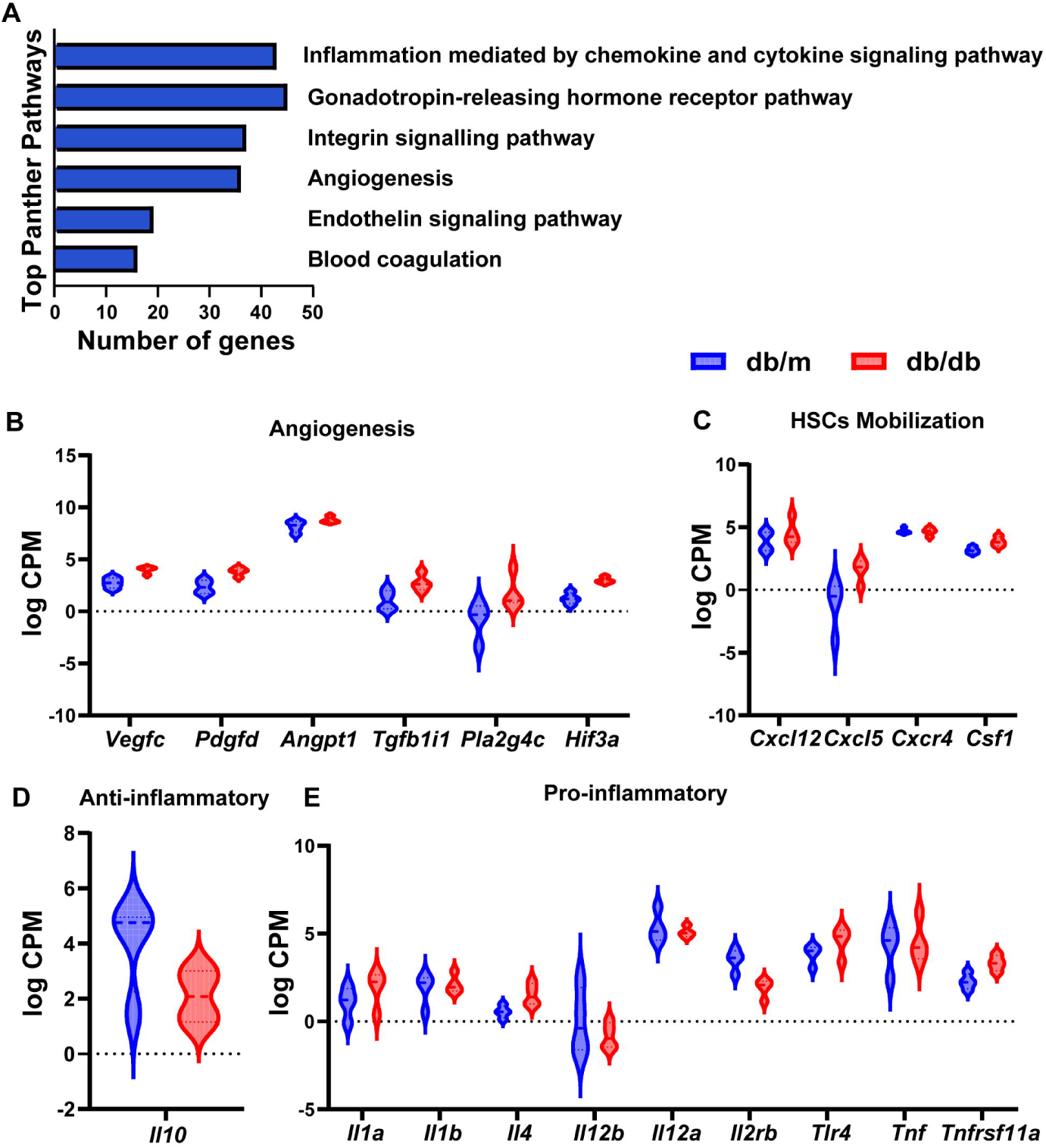
Transcriptome analysis revealed the gene level changes in the HSCs of db/db mice. The significant changes in mRNAs were analyzed using Gene ontology/Panther to uncover the top pathways. **A)** The pathways with the highest number of differentially expressed genes in the HSCs of db/db and db/m mice. Individual pathways analysis showed increased expression of **B)** angiogenesis, **C)** differentiation, and migration of HSCs-related genes. **D)** The anti-inflammatory gene IL-10 was significantly downregulated in the db/db mice, **E)** whereas there was an increase in the expression level of pro-inflammatory genes in db/db mice.

### Overexpression of inflammatory pathways in the HSCs of db/db mice

For further pathway analysis, we chose 446 genes which were also present in our mRNA sequencing database. Interestingly, out of 446 genes, 90 were targeted by multiple miRNAs (**Figure 3**). These target mRNAs could be of more potential as their regulation is controlled by more than one miRNA. We ran a core pathway analysis on significantly changed miRNAs to investigate the potential pathways associated with differentially expressed miRNAs. The top 5 miRNAs (both upregulated and downregulated) (**Table S1**) were further subjected to the network analysis and overlaid with the mRNA from our sequencing database. Since our group has previously shown the role of miR-92 in diabetic retinopathy (Luo et al., 2022, 2023), we also included miR-92 (fold change -1.577 and p=0.0065) for the network analysis. Lastly, these top miRNAs were subjected to the IPA pathway analysis.

**Figure 3.**
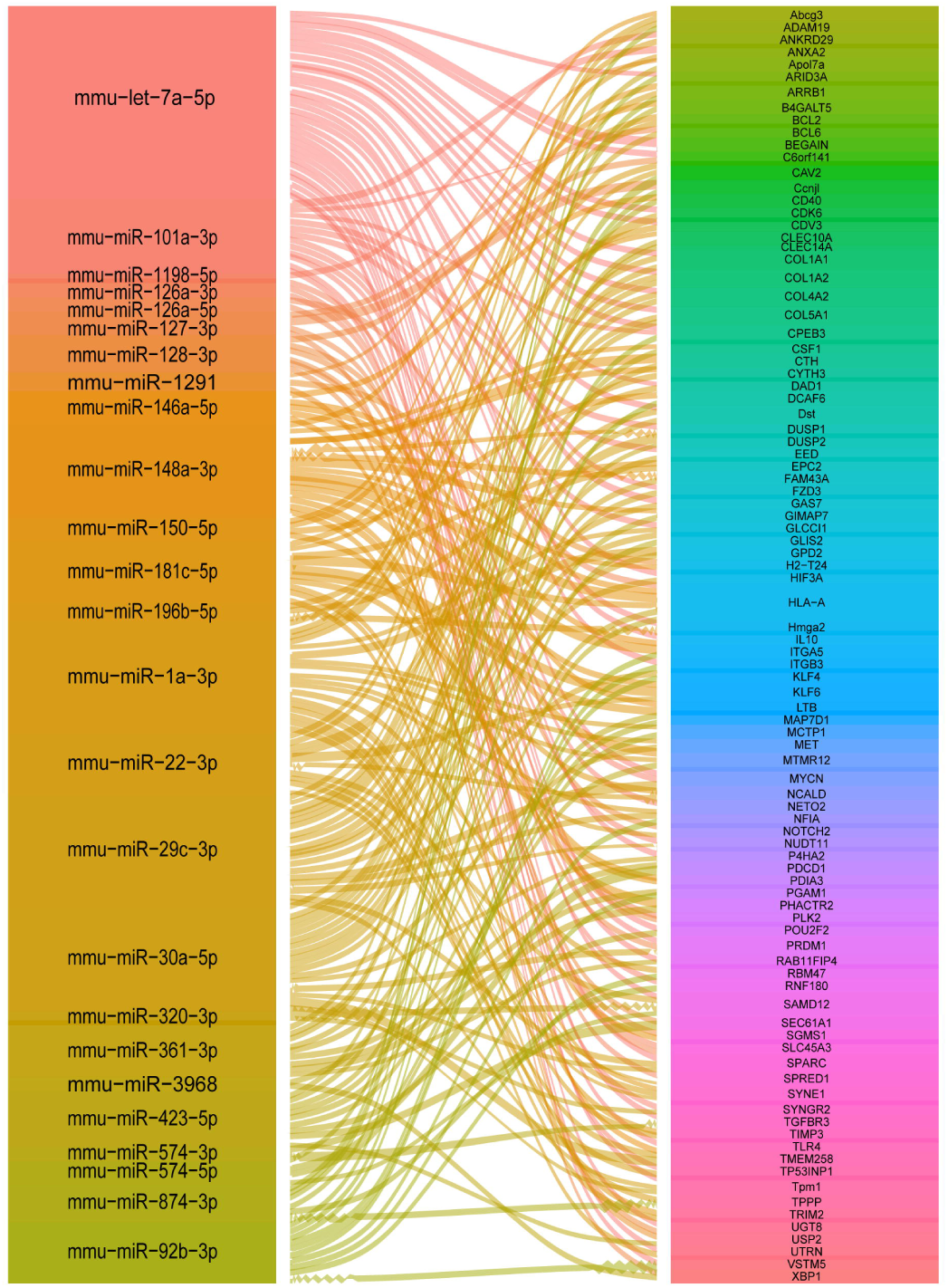
Alluvial plot illustrating overlapped mRNA targets between multiple miRNAs. MicroRNA target filter analysis revealed a total of 1974 mRNA targets, which were then overlaid with the significantly expressed mRNA targets. 446 mRNA targets were common among two datasets from which 206 mRNAs were targeted by multiple miRNAs, as shown in the Figure. This Figure was plotted using a freely available online tool SRplot (https://www.bioinformatics.com.cn/en).

IPA pathway and network analysis showed that each miRNA was targeting inflammation-related pathways. miRNAs are a known transcriptional regulator, where they can bind to the 3’-UTR regions of the mRNA leading to their degradation. With the differential expression of the top regulated miRNAs [miR-874-3p, miR-30c-5p, miR-3968 and miR-148a-3p **(Figure 4A-D);** and miR-1-3p, miR-423-5p, miR-22-3p, miR-574-5p, and miR-92a-3p (**Figure S2-S4**) of the LSK cells, we found an increase in the expression of downstream targets like colony-stimulating factor 1 (CSF1), Kruppel-like factors (*Klf6, Klf4*, and *Klf2*), integrins (*Itga5, Itga9, Itgav, Itgb3*), and collagen (*Col1a1, Col1a2, Col5a1, Col4a2*) all of which held the potential to stimulate the chemotaxis, differentiation and proliferation of bone-marrow-derived monocytes/macrophages and microglia. The top canonical pathways affected by these miRNAs and their respective targets turned out to be macrophage alternative activation signalling, neuroinflammation, IL-4 signalling, TGF-β signalling, CCR5 signalling in macrophages, leukocyte extravasation signalling, VEGF family ligand-receptor interactions, neurovascular coupling signalling, neutrophil extracellular trap signalling, phagosome formation, WNT/Ca^+^ signalling, IL-12 signalling, natural killer cell signalling and CREB signalling in neurons (**Figure 4A** and **Figure S2,3,4**). The IPA suggested a shift to the pro-inflammatory phenotype of HSCs in the db/db mice at the transcriptional level, which in turn is largely regulated by the miRNAs.

**Figure 4.**
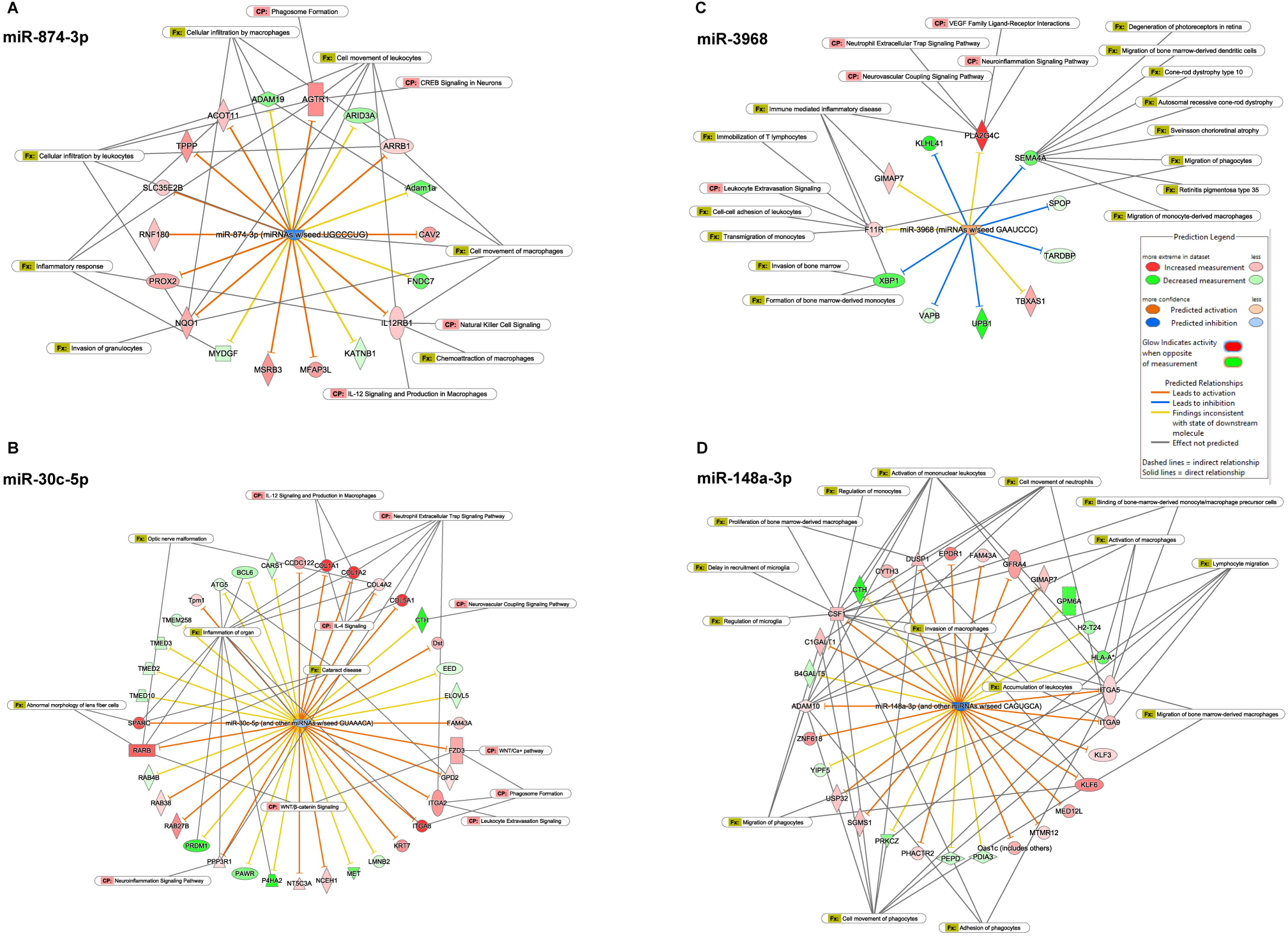
Top pathway analysis for miRNAs reflected increased inflammation-related genes in db/db mice. Top miRNAs were analyzed using IPA software for their downstream targets and associated pathways. **A)** miR-874-3p and **B)** miR-30c-5p were upregulated in the HSCs of db/db mice, which were traced back to leukocyte extravasation, neuroinflammation, cytokine signaling, and macrophage signaling. **C)** and **D)** miR-3968 and miR-148a-3p were top downregulated miRNAs, targeting VEGF signaling, formation, proliferation, and migration of bone marrow-derived macrophages and monocytes, recruitment, and migration of microglia, neuroinflammation and degeneration of photoreceptors. The data was uploaded to the IPA, and miRNAs and mRNAs with a p<0.05 were used for the miRNA target filter and core pathway analysis.

### Chronic diabetes accelerates retinal inflammation

One of the devastating consequences of diabetes is enhanced inflammation, which significantly contributes to the associated vascular complications. To study this, the retinas were isolated and stained with cell surface markers corresponding to the local and peripheral inflammatory cell types to assess the retinal inflammation in these animals. Using flow cytometry, we quantified the total microglia, resident microglia, and monocyte populations. **Figure 5A** shows a representative gating strategy used for the flow cytometric analysis. We found a significant increase in the total microglia population [represented as CD11b^high^; CD45^low^; Ly6G^-^; Ly6C^-^; F4/80^mid^] in the db/db mice (**Figure 5B**), however further gating on this population for CD39 showed a decrease in the db/db animals (**Figure 5C**). CD39 is a marker for the resident microglia (Hu et al., 2017), a reduction in the CD39^+^ microglia depicts an increase in the infiltrating inflammatory microglia from the peripheral system. The quantification for monocytes (CD11b^high^; CD45^high^; Ly6G^+^; Ly6C^+^; F4/80^mid^) showed an increase in the db/db mice (**Figure S5A**), however, the change was not significant.

**Figure 5.**
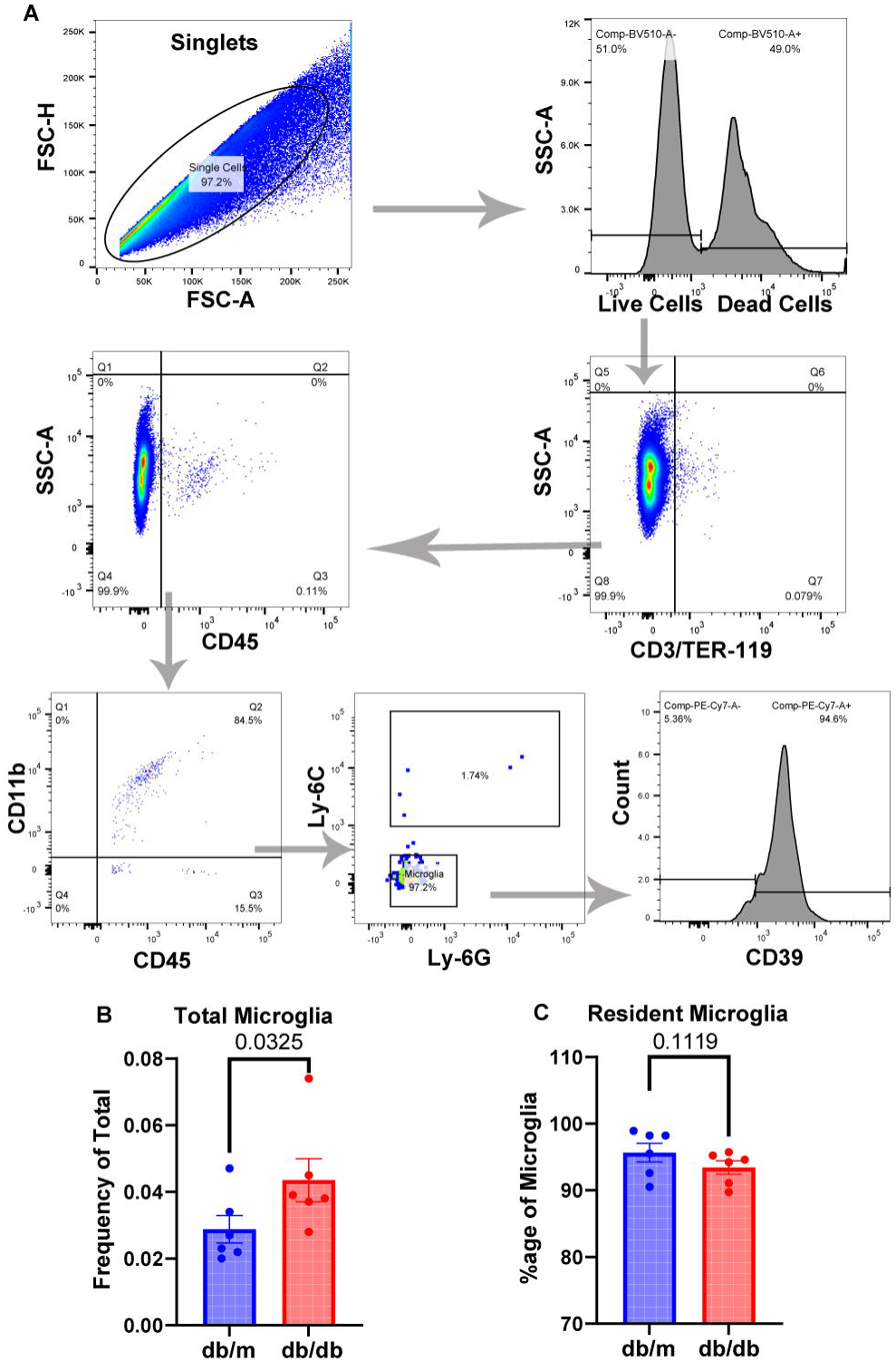
Chronic diabetes enhances retinal inflammation in db/db mice. Flow cytometric analysis was conducted to quantify the inflammatory microglia population in the retinas of db/db mice. **A)** The representative gating strategy used for flow cytometric analysis. The microglial population was gated as CD11b^+^CD45^mid^LyC^-^LyG^-^. The microglia were further gated for CD39, a cell surface marker for resident microglia. **B)** The microglia population was increased in the db/db mice. **C)** There was a noticeable decline in the resident microglia (CD11b^+^CD45^mid^LyC^-^ LyG^-^ CD39^+^) in db/db mice. The data were presented as mean ± SEM; N=6 and analyzed using a student’s t-test.

### Chronic diabetes impairs retinal structure and delays retinal function

Since chronic diabetes is also associated with the neuro-vascular deficits of the retina, we also examined the structural and functional damages using OCT (**Figure 6A**), FA (**Figure 6E**), and ERG (**Figure 6H-O**). The OCT data analysis showed a significant reduction in the total retinal thickness in the db/db mice compared to their age-matched db/m control mice (**Figure 6B)**. Also, the outer nuclear layer (ONL) thickness was found to be decreased in db/db animals (**Figure 6D**), with no significant changes in the inner nuclear layer (INL) thickness (**Figure 6C**). Further, to assess the vascular defects in the retinas of diabetic mice, the Fluorescein Angiography (FA) was performed. Fluorescein dye was injected intraperitoneally into the animals, followed by fundoscopy (**Figure 6E**). Fundus images analysis using MATLAB software showed a higher vein tortuosity in db/db mice, with no significant change in the vessel width (**Figure 6F, G**). Altogether, OCT and FA showed diabetes-induced deficits in the retinal structure and vasculature in db/db mice.

**Figure 6.**
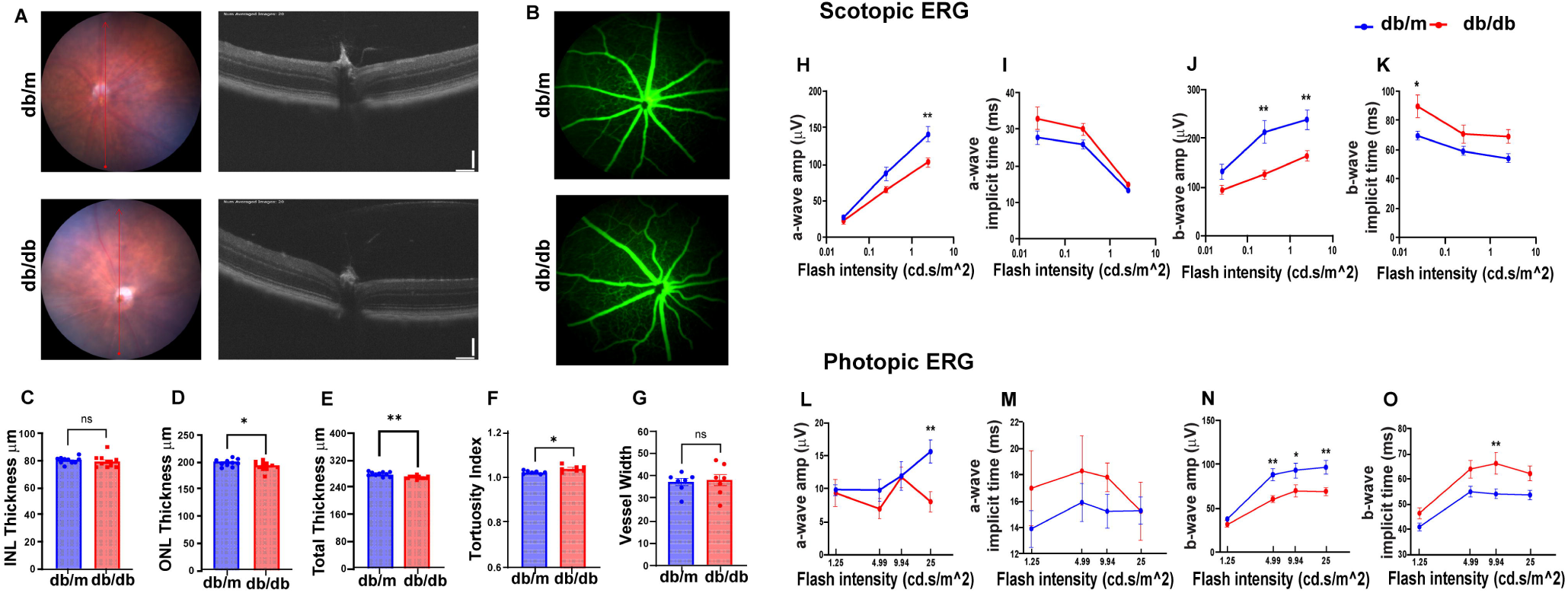
Chronic diabetes impairs retinal neuro-vasculature and function. 6-month-old diabetic mice’s retinas were analyzed for structural abnormalities using OCT and FA. **A)** Representative fundus and retinal layers images captured using OCT. **(B-D)** Total retinal thickness, inner nuclear layer, and outer nuclear layer thickness quantified using in-built software provided by Phoenix Micron IV showed a decrease of the total and ONL in db/db mice. **E)** Representative fundus images captured after i.p administration of fluorescein. Diabetes increased **(F)** vessel tortuosity, with no significant change in the vessel width **(G)**. Db/db mice showed reduced scotopic ERG a-wave **(H)** and b-wave **(J)** responses when compared to their age-matched db/m control mice. The implicit time for a-wave **(I)** and b-wave **(K)** was higher in db/db mice. The photopic a-wave amplitude **(L)**, implicit time **(M)**; b-wave amplitude, and implicit time **(N & O)** had a similar pattern as of scotopic ERG in the db/db mice. Data are expressed as mean ± SEM. **p<0.01; *p<0.01 show changes vs db/m mice; Scale bar-100 µM. N=9 for A-D; N=7 for E-G and N=10 for H-K, statistical test used is a student t-test for A-G and the data were analyzed using two-way ANOVA followed by Sidak’s multiple comparisons test for H-O. INL-inner nuclear layer; ONL-outer nuclear layer.

Since we observed defects in the retinal structure, the next retinal function was assessed in these mice using ERG. The a-wave originating from the photoreceptors and the b-wave originating from bipolar cells were analyzed under scotopic (dark-adapted; rods-driven) and photopic (light-adapted) conditions. In both scotopic and photopic conditions, the a-wave amplitude was decreased in the db/db mice at higher flash intensity, suggesting a decrease in the retinal photoreceptors’ activity (**Figure 6H**, and **6L**). Similarly, the b-wave amplitude was also reduced in the db/db mice irrespective of scotopic or photopic condition (**Figure 6J and 6N**), suggesting an impairment in bipolar cells in chronic diabetes. At the same time, we found a delay in the implicit time for both a-wave and b-wave in db/db mice. The a-wave implicit time was significantly higher in the db/db mice under photopic conditions; however, it was non-significant at individual flash intensities, and under scotopic conditions, the change was non-significant (**Figure 6I** and **6M**). The b-wave implicit time was also increased in the db/db mice under both conditions (**Figure 6K** and **6O**).

The oscillatory potential (OP) amplitude average in db/db mice was found to be decreased in the photopic ERG only (**Figure S5B**), with no significant change under scotopic ERG (**Figure S5C**). Overall, the ERG currents measurement and their quantification showed a distorted retinal function in the db/db mice compared to their age-matched control (db/m) mice.

## Discussion

In the present study, we have investigated the transcriptional dysfunctions of bone marrow-derived HSCs due to chronic diabetes, which have not been explored before. These transcriptional changes can, therefore, change the fate of circulatory HSCs. Through our transcriptional profiling, we identified a novel set of miRNAs (miR-3968, miR-574-5p, and miR-1971) and a distinguished set of mRNAs targeting angiogenesis and pro-inflammatory pathways in HSCs of diabetic animals with the potential to participate in microvascular complications such as retinopathy.

Bone marrow-derived HSCs are widely studied and used as therapeutics for malignant blood disorders due to their outstanding capacity to regenerate the hemato-lymphoid system (Epah and Schäfer, 2021). In the last two decades, multiple studies have reflected the negative outcome of diabetes on the HSCs, in terms of bone marrow function, release and migration ability of HSCs, and decreased circulation in the peripheral blood (Fadini et al., 2007). It is well-appreciated now that chronic diabetes can lead to an impaired endothelial repair system and promote stem cell mobilopathy and neuropathy (Bhatwadekar et al., 2017; Busik et al., 2009; Fadini and Albiero, 2022; Fadini et al., 2013). Consistent with the previous literature, db/db animals in the present study had a significantly higher number of entrapped HSCs in the bone marrow (Busik et al., 2009). Stem cell mobilization is largely dependent on the CXCL12-CXCR4 signalling, where the CXCL12 upregulation is linked to the impaired stem cell trafficking/mobilopathy in diabetes (Fadini et al., 2015; Ferraro et al., 2011). The higher expression of *Cxcl12,* with the simultaneous increase in the entrapped HSCs, reflected the inability of the HSCs to be released in diabetic animals.

The present study is the first of this kind, where the pure LSK cells were sorted and utilized to investigate diabetes-induced transcriptional changes in the db/db mice. Further, there is growing evidence that miRNAs play a crucial role in the transcriptional regulation of enormous molecular targets associated with metabolic disorders, including diabetes or diabetes-induced vascular complications (Ismail et al., 2023; Kolfschoten et al., 2009; Zampetaki and Mayr, 2012). However, it is noteworthy that most of these studies have been done on circulatory or tissue-specific miRNAs, and no data is available for the LSK miRNAs. Therefore, we investigated the LSK miRNA and mRNA targets due to their ability to change the fate of the stem cells being circulated and used for vascular repair. We found 35 significantly changed miRNAs in the LSK cells of db/db mice, with 25 being downregulated and 10 being upregulated. Interestingly, our sequencing data has uncovered 2 novel miRNAs (miR-3968, and miR-1971), which have never been reported before for any involvement with diabetes-induced microvascular complications or even diabetes alone. On the other hand, there were several miRNAs consistent with the reported literature, including miR-1-3p (Liu et al., 2020; Morales-Sánchez et al., 2023), miR-30c-5p (Dong and Wang, 2019; Mazzeo et al., 2018), miR-101-3p (Higuchi et al., 2015), miR-181a-5p (Cao et al., 2022), miR-340-5p (Zhu et al., 2022), miR-29b-3p (Zeng et al., 2020), miR-148a-3p (Liang et al., 2018) miR-423-5p (Blum et al., 2019) and miR-22-3p (Zhou et al., 2021). Contrarily, we found three miRNAs (miR-126a-5p and miR-126a-3p (Zampetaki et al., 2010); miR-874-3p (Li et al., 2021)) inconsistent with the reported miRNAs in terms of their expression pattern. Since this study is distinguishable from the available literature due to the use of HSCs, the miRNAs observed in the data hold great potential irrespective of their expression difference with the previous studies.

The miRNAs target several downstream pathways that can manifest the fate of HSCs, to explore this phenomenon, we further analyzed the top miRNAs. We found multiple miRNAs and their downstream mRNAs targeting inflammatory and angiogenic pathways. One such target is Kruppel-like factors (KLFs); a family of transcription factors that regulate macrophage differentiation and migration under the influence of inflammatory signals (Cao et al., 2010). Interestingly, there was a significant increase in the expression levels of *Klf4* and K*lf6* in the LSK cells of diabetic animals, possibly contributing to the migration of inflammatory monocytes/macrophages (Alder et al., 2008; Date et al., 2014; Feinberg et al., 2007). Moreover, we also found a decrease in the expression of miR-92a-3p in the LSK cells, which was previously shown to decrease in the angiogenic cells of diabetic retinopathy individuals (Bhatwadekar et al.) and retinas of diabetic mice (Kovacs et al., 2011). Also, it has been shown that interactions of miR-92a-3p with KLF2 and KLF4 can modulate inflammatory macrophage signaling (Fang and Davies, 2012; Georgantas et al., 2007), thus highlighting the possible involvement of miR-92a-3p regulated inflammatory pathways in diabetic HSCs. Likewise, studies have shown the role of colony-stimulating factor 1 (CSF1) in microglia activation and inflammatory cytokine secretion (Kokona et al., 2018). We here found a significantly higher expression of the *Csf1* gene in diabetic animals, thus promoting microglia activation and inflammation. A higher expression of pro-inflammatory genes *Tlr4, Il1*α, and *Tnf11*α and a decrease in the expression levels of anti-inflammatory cytokine *Il10*, in the LSK cells, also suggested a shift to inflammatory phenotype (D’alessandra et al., 2021; Yue et al., 2022) in diabetic HSCs. Moreover, there was a higher expression of the *Ccl4* gene, which regulates macrophage activation, chemotaxis, and migration. A simultaneous increase of these genes in the LSK cells of db/db mice marks the initiation of inflammation at transcriptional levels in the stem cells.

In addition to the changes in the inflammatory genes, we found several angiogenic genes being upregulated in the LSK cells of db/db mice. Vascular endothelial growth factor (VEGF) is a widely studied angiogenic factor with multiple isoforms (Melincovici et al., 2018). VEGF-C, primarily known for lymphangiogenesis, has also been reported to play a part in pathological angiogenesis (Nagai and Minami, 2015). An increase in the gene expression of *Vegfc* in db/db mice reflects the angiogenic fate of HSCs.

Numerous studies have shown that chronic diabetes severely deteriorates the retinal health of diabetic individuals and experimental animal models. In accordance with the previous literature, we also found that db/db animals had decreased retinal thickness (Yang et al., 2015), their overall retinal function was compromised (Bogdanov et al., 2014), and more tortuosity was observed in the retinal blood vessels of db/db mice (Bhatta et al., 2015). Our study showed a defect in the photoreceptors and bipolar cells, as evident from the decreased currents during scotopic and photopic ERG. These defects in the retina were reflected in the infiltration of microglial cells, suggesting an essential role of bone marrow-derived cells in structural and functional abnormalities of diabetes (Altmann and Schmidt, 2018; Asare-Bediako et al., 2022; Chakravarthy et al., 2016). Indeed, we observed an increase in the total microglia population (CD11b^high^; CD45^low^; Ly6G^-^; Ly6C^-^; F4/80^mid^] in the db/db mice. Consistent with previous studies (Fu et al., 2023; Hu et al., 2017), while there was a decrease in resident microglial populations, this difference was not statistically significant. We speculate this may be due to the duration of diabetes, analysis type (immunofluorescence vs. flow cytometry), and sample size.

In summary, we have observed HSCs mobilopathy, accompanied by adverse effects on the transcriptome profile of LSK cells in chronically diabetic animals. Several miRNAs and mRNA targets reflected a shift to inflammatory and angiogenic fate of bone-marrow derived HSCs in diabetes. The investigation of HSCs transcriptome is clinically very relevant as these cells not only participate in the pathogenesis of disease but also play an important role in hematopoiesis and vascular repair. We believe that the present study has strengthened the current knowledge of diabetes-induced HSC dysfunction at the transcriptional levels, with the potential to contribute to vascular damage. Identifying novel miRNAs and their targets could be a potential biomarker for diabetes-induced vasculature deficits such as retinopathy. However, further studies are required to investigate the molecular mechanisms linking the HSCs miRNAs or mRNA targets to the progression of vascular disease in diabetes.

## Experimental Procedures

### Resource Availability

Corresponding author: Further information and requests for resources and reagents should be directed to and will be fulfilled by the lead corresponding author, Ashay Bhatwadekar.

#### Material availability

This study did not generate new unique reagents.

#### Data and code availability

The sequencing data will be uploaded online after the manuscript is accepted.

### Animals

The B6.BKS(D)-Lepr^db^/J (an animal model for T2D; db/db) and Lepr^db^/^+^ db/m (heterozygotes; db/m) mice [stock number 000697] were procured from The Jackson Laboratory (Bar Harbor, ME, USA) and housed in the animal care facility at Glick eye institute, Indiana university. All the animals were kept under normal physiological conditions (12-hour light/dark conditions), with free access to food and water all the time. The mice were maintained for eight months (6 months of diabetes) before proceeding with the retinal exams and related experiments.

All the experiments performed were per the Guiding Principles in the Care and Use of Animals (National Institutes of Health) and the Association for Research in Vision and Ophthalmology’s Statement for the Use of Animals in Ophthalmic and Vision Research.

### Electroretinogram (ERG)

The ERG was performed to assess the retinal function under both scotopic (dark-adapted) and photopic (light-adapted) conditions. The mice were dark-adapted overnight before the day of the experiment. The mice were anesthetized with an i.p injection of ketamine/xylazine mixture, followed by the topical application of 1% tropicamide/2.5% phenylephrine to dilate pupils. To keep the eyes moist during the measurements, Gonak® or Hypromellose 2.5% solution (Akorn, Lake Forest, Illinois, USA) was applied. The reference electrode and the ground needle were placed sub dermally between the eyes and at the base of the tail, respectively. The gold loop electrodes were then placed over the corneas. Lastly, the recordings were captured using a LKC NGIT-100 recording machine (LKC Technologies, Inc, Gaithersburg, MD, USA). For photopic ERG, mice were light-adapted for 10 min, followed by the scotopic recordings. The a-wave, b-wave amplitude, Oscillatory potentials, and implicit times were calculated using in-built analysis tools from LKC Technologies.

### Optical Coherence Tomography (OCT)

Retinal thickness was measured using the Micron IV OCT System (Phoenix Technology Group, Pleasanton, CA, USA). The mice were anesthetized similarly as described above in ERG, followed by pupil dilation. The optic nerve was located manually and used as a center for the scans. Both eyes were scanned, and retinal thickness was measured using in-built software provided by the Micron IV OCT system.

### Fluorescein Angiography (FA)

The mice were administered intraperitoneally with fluorescein (@ 5ml/kg of 2.5% fluorescein solution) after each mouse was anesthetized using the same method as described in ERG and OCT. FA was then performed on the Micron IV OCT system, and images were analyzed using MATLAB software.

### Microglia analysis

The eyes were collected in PBS after animal sacrifice. The retinal samples for staining were prepared according to the method described previously (Wu et al., 2023). Briefly, the retinas were isolated and incubated in the digestion buffer for 30 min at 37°C in a water bath. Following the incubation, the tissues were disrupted using a 10 ml syringe, washed with FACS buffer, and centrifuged at 400g for 5 min at 4°C. The supernatant was discarded, and 1 ml of FACS buffer was added to the cell pellet, followed by staining with the cell surface marker antibodies. The single cell suspensions of retinas were stained with Zombie Aqua^TM^ Fixable Viability Kit (BioLegend) for 20 min and then washed with PBS. Cells were incubated with a combination of fluorophore-conjugated primary antibodies against mouse CD3, CD11b, CD45, TER-119, F4/80, Ly-6G, Ly-6C, and CD39 (BioLegend, San Diego, CA, USA), for 30 min at room temperature. After staining, cells were washed and fixed with 2% paraformaldehyde in PBS. Finally, data were acquired on BD LSR Fortessa X-20 Flow cytometer using BD FACSDiva software (BD Biosciences, San Jose, CA, USA) and analyzed on FlowJo software by BD Biosciences (https://www.flowjo.com/).

### Hematopoietic stem cell (HSCs) isolation

Femur and tibia were collected for bone marrow isolation after animal sacrifice. Briefly, all the surrounding muscles were removed, and bones were cleaned and flushed with FACS buffer (PBS with 2% FBS) with the help of a needle. A-cell suspension was generated by triturating the cells through the needles. Cells were then treated with ice-cold ammonium chloride for 10 min to lyse all the red blood cells, followed by washing with PBS and staining with an anti-mouse lineage cocktail, Sca1, and c-Kit/CD117 antibodies (STEMCELL Technologies) for 30 min. The samples were finally sorted for lin^-^Sca1-ckit^+^ cells using a flow cytometer. The pure population of lin^-^Sca1-ckit^+^ cells represents HSCs used for the sequencing experiments.

### RNA isolation and sequencing

The RNA was extracted from the sorted HSCs using the Trizol-chloroform method followed by DNase treatment in the solution and cleaned up with an RNeasy MinElute clean-up kit from Qiagen. The purity of RNA was confirmed on the Bioanalyzer before proceeding with the sequencing. Only the samples with a RIN (RNA integrity number) of > 7 were used for the sequencing. The samples were prepared at the medical genomics core facility (Indiana University) for the paired-end mRNA and single-end miRNA seq library and sequenced on Illumina NovaSeq 6000 with a read length of 100bp for mRNA and NextSeq 2000 with a read length of 75 bp. The mRNA sequence reads were mapped to the designated reference genome using STAR (Spliced Transcripts Alignment to a Reference). The miRNA sequence reads were mapped to the designated reference genome using the Qiagen GeneGlobe RNA-seq Analysis Portal with default parameters for miRNA. The differential gene expression analysis for both mRNA and miRNA was performed with edgeR (Robinson, Mccarthy, and Smyth 2010). In this workflow, the statistical methodology uses negative binomial generalized linear models with likelihood ratio tests.

### Pathway Analysis

Ingenuity pathway analysis (IPA) software (QIAGEN Inc., https://www.qiagenbioinformatics.com/products/ingenuityLJpathwayLJanalysis) was used to identify biological/molecular pathways associated with the differentially expressed miRNAs and their targets. Both miRNAs and mRNA data were uploaded to the IPA and were subjected to the threshold filter of p-value -0.05. Then, a Micro RNA target filter was applied to find the mRNA targets for the differentially expressed miRNAs. The filter selections for our analysis included p<0.05, experimentally observed interactions, and direct and indirect relationships. mRNA targets from the IPA were overlaid with the mRNA targets from our sequencing database, and finally, pathways analysis was done for those miRNAs and mRNAs. mRNAs were also subjected to Gene Ontology and Panther analysis (pantherdb.org).

### Statistical Analysis

All the data were expressed as Mean+ SEM and analyzed using GraphPad Prism 10.0.1 for Windows (San Diego, California; www.graphpad.com). The intergroup comparison was done using a t-test and was considered significant when the p-value was less than 0.05. The ERG data was analyzed using two-way ANOVA followed by Sidak’s multiple comparisons test.

## Acknowledgments

The authors would like to acknowledge the funding support from the National Institute of Health (NIH)—National Eye Institute (NEI) grants, R01EY027779 R01EY027779-S1 and R01EY032080 to AB. Additionally, the Department of Ophthalmology is supported by an Unrestricted Grant from Research to Prevent Blindness (RPB). We would also like to thank flow cytometry and the Centre for Medical Genomics (CMG) core of Indiana University for helping us with the cell sorting and sequencing experiments. We want to thank Dr. Daniel Saban, Department of Ophthalmology, Duke University, for the helpful discussion on microglial analysis of the retina.

## Authors’ Contributions

Conceptualization: AB, NM; Validation: NM, AB; Formal Analysis: NM, AB; Investigation: NM, QL; Resources: AB; Data Curation: NM, QL, SA; Writing-Original Draft: NM; Writing-Review & Editing: AB, NM; Visualization: NM, AB; Supervision: QL, AB; Project Administration: AB, QL, NM; Funding Acquisition: AB.

---

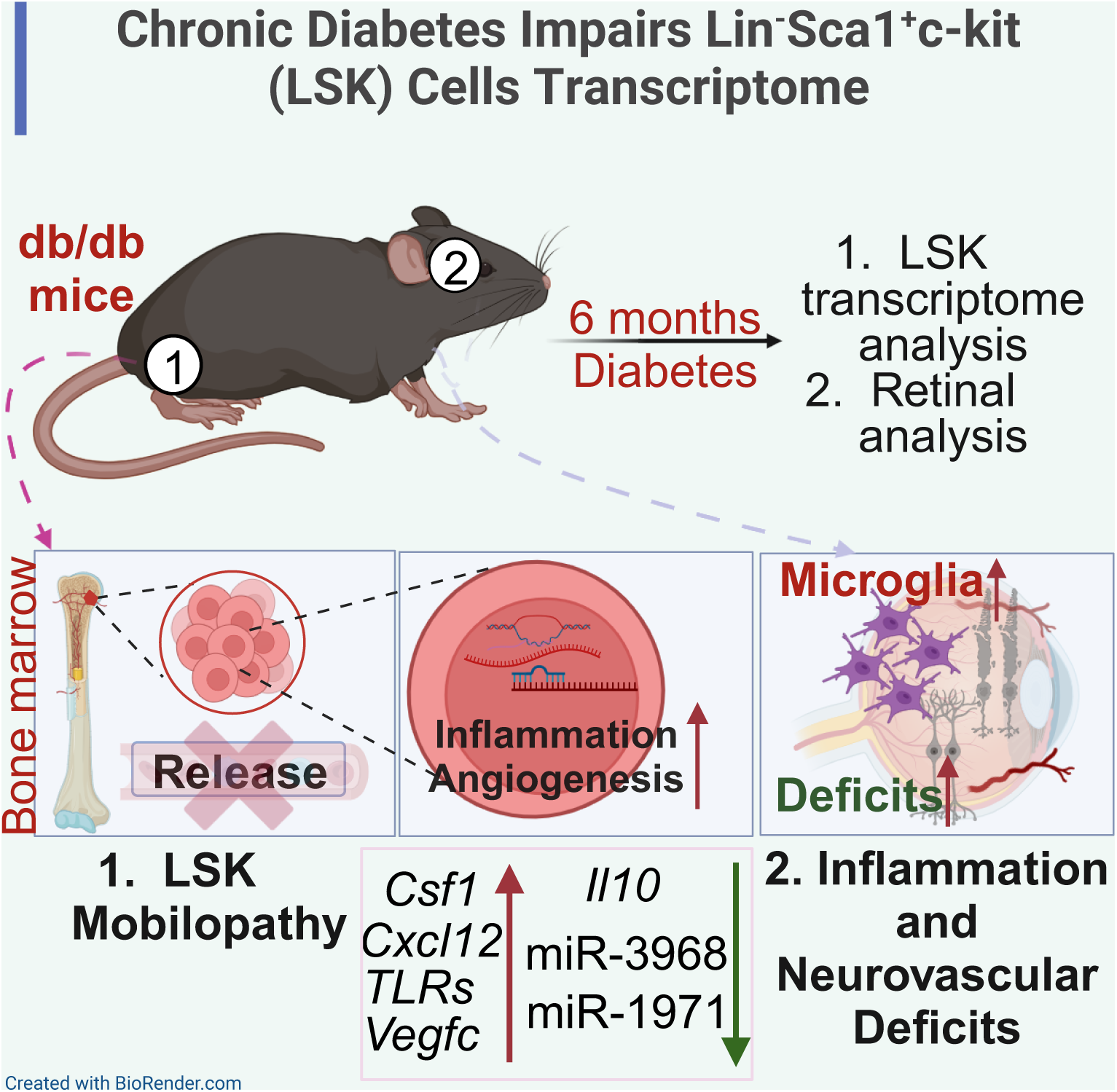

## References

Albiero, M., Bonora, B.M., and Fadini, G.P. (2020). Diabetes pharmacotherapy and circulating stem/progenitor cells. State of the art and evidence gaps. Curr. Opin. Pharmacol. 55. 10.1016/j.coph.2020.10.019.

Alder, J.K., Georgantas, R.W., Hildreth, R.L., Kaplan, I.M., Morisot, S., Yu, X., McDevitt, M., and Civin, C.I. (2008). Kruppel-Like Factor 4 Is Essential for Inflammatory Monocyte Differentiation In Vivo. J. Immunol. 180. 10.4049/jimmunol.180.8.5645.

Altmann, C., and Schmidt, M.H.H. (2018). The role of microglia in diabetic retinopathy: Inflammation, microvasculature defects and neurodegeneration. Int. J. Mol. Sci. 19. 10.3390/ijms19010110.

Asare-Bediako, B., Adu-Agyeiwaah, Y., Abad, A., Li Calzi, S., Floyd, J.L., Prasad, R., DuPont, M., Asare-Bediako, R., Bustelo, X.R., and Grant, M.B. (2022). Hematopoietic Cells Influence Vascular Development in the Retina. Cells 11. 10.3390/cells11203207.

Bhatta, M., Ma, J.H., Wang, J.J., Sakowski, J., and Zhang, S.X. (2015). Enhanced endoplasmic reticulum stress in bone marrow angiogenic progenitor cells in a mouse model of long-term experimental type 2 diabetes. Diabetologia 58. 10.1007/s00125-015-3643-3.

Bhatwadekar, A., Yan, Y., Stepps, V., Diabetes, S.H.-, and 2015, undefined miR-92a Corrects CD34+ Cell Dysfunction in Diabetes by Modulating Core Circadian Genes Involved in Progenitor Differentiation. Am Diabetes Assoc.

Bhatwadekar, A.D., Duan, Y., Korah, M., Thinschmidt, J.S., Hu, P., Leley, S.P., Caballero, S., Shaw, L., Busik, J., and Grant, M.B. (2017). Hematopoietic stem/progenitor involvement in retinal microvascular repair during diabetes: Implications for bone marrow rejuvenation. Vision Res. 139. 10.1016/j.visres.2017.06.016.

Blum, A., Meerson, A., Rohana, H., Jabaly, H., Nahul, N., Celesh, D., Romanenko, O., and Tamir, S. (2019). MicroRNA-423 may regulate diabetic vasculopathy. Clin. Exp. Med. 19. 10.1007/s10238-019-00573-8.

Bogdanov, P., Corraliza, L., Villena, J.A., Carvalho, A.R., Garcia-Arumí, J., Ramos, D., Ruberte, J., Simó, R., and Hernández, C. (2014). The db/db mouse: A useful model for the study of diabetic retinal neurodegeneration. PLoS One 9. 10.1371/journal.pone.0097302.

Busik, J. V., Tikhonenko, M., Bhatwadekar, A., Opreanu, M., Yakubova, N., Caballero, S., Player, D., Nakagawa, T., Afzal, A., Kielczewski, J., et al. (2009). Diabetic retinopathy is associated with bone marrow neuropathy and a depressed peripheral clock. J. Exp. Med. 206. 10.1084/jem.20090889.

Cao, J., Zhao, C., Gong, L., Cheng, X., Yang, J., Zhu, M., and Lv, X. (2022). MiR-181 Enhances Proliferative and Migratory Potentials of Retinal Endothelial Cells in Diabetic Retinopathy by Targeting KLF6. Curr. Eye Res. 47. 10.1080/02713683.2022.2039206.

Cao, Z., Sun, X., Icli, B., Wara, A.K., and Feinberg, M.W. (2010). Role of Krüppel-like factors in leukocyte development, function, and disease. Blood 116. 10.1182/blood-2010-05-285353.

Chakravarthy, H., Beli, E., Navitskaya, S., O’Reilly, S., Wang, Q., Kady, N., Huang, C., Grant, M.B., and Busik, J. V. (2016). Imbalances in mobilization and activation of pro-inflammatory and vascular reparative bone marrow-derived cells in diabetic retinopathy. PLoS One 11. 10.1371/journal.pone.0146829.

Corbi, S.C.T., de Vasconcellos, J.F., Bastos, A.S., Bussaneli, D.G., da Silva, B.R., Santos, R.A., Takahashi, C.S., de S. Rocha, C., Carvalho, B. de S., Maurer-Morelli, C. V., et al. (2020). Circulating lymphocytes and monocytes transcriptomic analysis of patients with type 2 diabetes mellitus, dyslipidemia and periodontitis. Sci. Rep. 10. 10.1038/s41598-020-65042-9.

D’alessandra, Y., Chiesa, M., Vigorelli, V., Ricci, V., Rurali, E., Raucci, A., Colombo, G.I., Pompilio, G., and Vinci, M.C. (2021). Diabetes induces a transcriptional signature in bone marrow– derived CD34+ hematopoietic stem cells predictive of their progeny dysfunction. Int. J. Mol. Sci. 22. 10.3390/ijms22031423.

Date, D., Das, R., Narla, G., Simon, D.I., Jain, M.K., and Mahabeleshwar, G.H. (2014). Kruppel-like transcription factor 6 regulates inflammatory macrophage polarization. J. Biol. Chem. 289. 10.1074/jbc.M113.526749.

Dong, N., and Wang, Y. (2019). MiR-30a Regulates S100A12-induced Retinal Microglial Activation and Inflammation by Targeting NLRP3. Curr. Eye Res. 44. 10.1080/02713683.2019.1632350.

Epah, J., and Schäfer, R. (2021). Implications of hematopoietic stem cells heterogeneity for gene therapies. Gene Ther. 28. 10.1038/s41434-021-00229-x.

Fadini, G.P., and Albiero, M. (2022). Impaired Hematopoietic Stem/Progenitor Cell Traffic and Multi-organ Damage in Diabetes. Stem Cells 40, 716–723. 10.1093/stmcls/sxac035.

Fadini, G.P., Sartore, S., Agostini, C., and Avogaro, A. (2007). Significance of endothelial progenitor cells in subjects with diabetes. Diabetes Care 30. 10.2337/dc06-2305.

Fadini, G.P., Albiero, M., De Kreutzenberg, S.V., Boscaro, E., Cappellari, R., Marescotti, M., Poncina, N., Agostini, C., and Avogaro, A. (2013). Diabetes impairs stem cell and proangiogenic cell mobilization in humans. Diabetes Care 36. 10.2337/dc12-1084.

Fadini, G.P., Fiala, M., Cappellari, R., Danna, M., Park, S., Poncina, N., Menegazzo, L., Albiero, M., Dipersio, J., Stockerl-Goldstein, K., et al. (2015). Diabetes limits stem cell mobilization following G-CSF but not plerixafor. Diabetes 64. 10.2337/db15-0077.

Fadini, G.P., Ciciliot, S., and Albiero, M. (2017). Concise Review: Perspectives and Clinical Implications of Bone Marrow and Circulating Stem Cell Defects in Diabetes. Stem Cells 35. 10.1002/stem.2445.

Fang, Y., and Davies, P.F. (2012). Site-specific microRNA-92a regulation of Krüppel-like factors 4 and 2 in atherosusceptible endothelium. Arterioscler. Thromb. Vasc. Biol. 32. 10.1161/ATVBAHA.111.244053.

Feinberg, M.W., Wara, A.K., Cao, Z., Lebedeva, M.A., Rosenbauer, F., Iwasaki, H., Hirai, H., Katz, J.P., Haspel, R.L., Gray, S., et al. (2007). The Kruppel-like factor KLF4 is a critical regulator of monocyte differentiation. EMBO J. 26. 10.1038/sj.emboj.7601824.

Ferraro, F., Lymperi, S., Méndez-Ferrer, S., Saez, B., Spencer, J.A., Yeap, B.Y., Masselli, E., Graiani, G., Prezioso, L., Rizzini, E.L., et al. (2011). Diabetes impairs hematopoietic stem cell mobilization by altering niche function. Sci. Transl. Med. 3. 10.1126/scitranslmed.3002191.

Fu, X., Feng, S., Qin, H., Yan, L., Zheng, C., and Yao, K. (2023). Microglia: The breakthrough to treat neovascularization and repair blood-retinal barrier in retinopathy. Front. Mol. Neurosci. 16. 10.3389/fnmol.2023.1100254.

Georgantas, R.W., Hildreth, R., Morisot, S., Alder, J., Liu, C.G., Heimfeld, S., Calin, G.A., Croce, C.M., and Civin, C.I. (2007). CD34+ hematopoietic stem-progenitor cell microRNA expression and function: A circuit diagram of differentiation control. Proc. Natl. Acad. Sci. U. S. A. 104. 10.1073/pnas.0610983104.

Hanoun, M., and Frenette, P.S. (2013). This Niche Is a Maze; An Amazing Niche. Cell Stem Cell 12, 391–392. 10.1016/J.STEM.2013.03.012.

He, X., Kuang, G., Wu, Y., and Ou, C. (2021). Emerging roles of exosomal miRNAs in diabetes mellitus. Clin. Transl. Med. 11. 10.1002/ctm2.468.

Higuchi, C., Nakatsuka, A., Eguchi, J., Teshigawara, S., Kanzaki, M., Katayama, A., Yamaguchi, S., Takahashi, N., Murakami, K., Ogawa, D., et al. (2015). Identification of circulating miR-101, miR-375 and miR-802 as biomarkers for type 2 diabetes. Metabolism. 64. 10.1016/j.metabol.2014.12.003.

Hu, P., Hunt, N.H., Arfuso, F., Shaw, L.C., Uddin, M.N., Zhu, M., Devasahayam, R., Adamson, S.J., Benson, V.L., Chan-Ling, T., et al. (2017). Increased indoleamine 2,3-dioxygenase and quinolinic acid expression in microglia and müller cells of diabetic human and rodent retina. Investig. Ophthalmol. Vis. Sci. 58, 5043–5055. 10.1167/iovs.17-21654.

IDF (2021). IDF Diabetes Atlas 2021 _ IDF Diabetes Atlas. IDF Off. Website 1–4..

Ismail, A., El-Mahdy, H.A., Eldeib, M.G., and Doghish, A.S. (2023). miRNAs as cornerstones in diabetic microvascular complications. Mol. Genet. Metab. 138. 10.1016/j.ymgme.2022.106978.

Kokona, D., Ebneter, A., Escher, P., and Zinkernagel, M.S. (2018). Colony-stimulating factor 1 receptor inhibition prevents disruption of the blood-retina barrier during chronic inflammation. J. Neuroinflammation 15. 10.1186/s12974-018-1373-4.

Kolfschoten, I.G.M., Roggli, E., Nesca, V., and Regazzi, R. (2009). Role and therapeutic potential of microRNAs in diabetes. Diabetes, Obes. Metab. 11. 10.1111/j.1463-1326.2009.01118.x.

Kovacs, B., Lumayag, S., Cowan, C., and Xu, S. (2011). microRNAs in early diabetic retinopathy in streptozotocin-induced diabetic rats. Investig. Ophthalmol. Vis. Sci. 52, 4402– 4409. 10.1167/iovs.10-6879.

Li, R., Yuan, H., Zhao, T., Yan, Y., Liu, Z., Cai, J., Qiu, C., and Li, C. (2021). miR-874 ameliorates retinopathy in diabetic rats by NF-κB signaling pathway. Adv. Clin. Exp. Med. 30. 10.17219/ACEM/130602.

Liang, Z., Gao, K.P., Wang, Y.X., Liu, Z.C., Tian, L., Yang, X.Z., Ding, J.Y., Wu, W.T., Yang, W.H., Li, Y.L., et al. (2018). RNA sequencing identified specific circulating mirna biomarkers for early detection of diabetes retinopathy. Am. J. Physiol. - Endocrinol. Metab. 315, E374– E385. 10.1152/AJPENDO.00021.2018.

Lin, Q., Zhou, W., Wang, Y., Huang, J., Hui, X., Zhou, Z., and Xiao, Y. (2020). Abnormal peripheral neutrophil transcriptome in newly diagnosed type 2 diabetes patients. J. Diabetes Res. 2020. 10.1155/2020/9519072.

Liu, J., Chen, S., Biswas, S., Nagrani, N., Chu, Y., Chakrabarti, S., and Feng, B. (2020). Glucose-induced oxidative stress and accelerated aging in endothelial cells are mediated by the depletion of mitochondrial SIRTs. Physiol. Rep. 8. 10.14814/phy2.14331.

Mazzeo, A., Beltramo, E., Lopatina, T., Gai, C., Trento, M., and Porta, M. (2018). Molecular and functional characterization of circulating extracellular vesicles from diabetic patients with and without retinopathy and healthy subjects. Exp. Eye Res. 176. 10.1016/j.exer.2018.07.003.

Melincovici, C.S., Boşca, A.B., Şuşman, S., Mărginean, M., Mihu, C., Istrate, M., Moldovan, I.M., Roman, A.L., and Mihu, C.M. (2018). Vascular endothelial growth factor (VEGF) – key factor in normal and pathological angiogenesis. Rom. J. Morphol. Embryol. 59..

Morales-Sánchez, P., Lambert, C., Ares-Blanco, J., Suárez-Gutiérrez, L., Villa-Fernández, E., Garcia, A.V., García-Villarino, M., Tejedor, J.R., Fraga, M.F., Torre, E.M., et al. (2023). Circulating miRNA expression in long-standing type 1 diabetes mellitus. Sci. Rep. 13. 10.1038/s41598-023-35836-8.

Nagai, N., and Minami, T. (2015). Emerging Role of VEGFC in Pathological Angiogenesis. EBioMedicine 2. 10.1016/j.ebiom.2015.11.006.

Paul, S., Ali, A., and Katare, R. (2020). Molecular complexities underlying the vascular complications of diabetes mellitus – A comprehensive review. J. Diabetes Complications 34, 107613. 10.1016/j.jdiacomp.2020.107613.

Smit-McBride, Z., and Morse, L.S. (2021). MicroRNA and diabetic retinopathy—biomarkers and novel therapeutics. Ann. Transl. Med. 9, 1280–1280. 10.21037/atm-20-5189.

Sun, H., Saeedi, P., Karuranga, S., Pinkepank, M., Ogurtsova, K., Duncan, B.B., Stein, C., Basit, A., Chan, J.C.N., Mbanya, J.C., et al. (2022). IDF Diabetes Atlas: Global, regional and country-level diabetes prevalence estimates for 2021 and projections for 2045. Diabetes Res. Clin. Pract. 183. 10.1016/j.diabres.2021.109119.

Tonyan, Z.N., Nasykhova, Y.A., Danilova, M.M., Barbitoff, Y.A., Changalidi, A.I., Mikhailova, A.A., and Glotov, A.S. (2022). Overview of Transcriptomic Research on Type 2 Diabetes: Challenges and Perspectives. Genes (Basel). 13. 10.3390/genes13071176.

Vinci, M.C., Gambini, E., Bassetti, B., Genovese, S., and Pompilio, G. (2020). When good guys turn bad: Bone Marrow’s and hematopoietic stem cells’ role in the pathobiology of diabetic complications. Int. J. Mol. Sci. 21, 1–22. 10.3390/ijms21113864.

Wu, T., Pelus, L.M., Artur Plett, P., Sampson, C.H., Chua, H.L., Fisher, A., Feng, H., Liu, L., Li, H., Ortiz, M., et al. (2023). Hematopoietic Acute Radiation Syndrome. 10.1667/RADE-22-00208.1 199, 468–489. https://doi.org/10.1667/RADE-22-00208.1.

Yang, Q., Xu, Y., Xie, P., Cheng, H., Song, Q., Su, T., Yuan, S., and Liu, Q. (2015). Retinal neurodegeneration in db/db mice at the early period of diabetes. J. Ophthalmol. 2015. 10.1155/2015/757412.

Yue, T., Shi, Y., Luo, S., Weng, J., Wu, Y., and Zheng, X. (2022). The role of inflammation in immune system of diabetic retinopathy: Molecular mechanisms, pathogenetic role and therapeutic implications. Front. Immunol. 13. 10.3389/FIMMU.2022.1055087/FULL.

Zampetaki, A., and Mayr, M. (2012). MicroRNAs in vascular and metabolic disease. Circ. Res. 110. 10.1161/CIRCRESAHA.111.247445.

Zampetaki, A., Kiechl, S., Drozdov, I., Willeit, P., Mayr, U., Prokopi, M., Mayr, A., Weger, S., Oberhollenzer, F., Bonora, E., et al. (2010). Plasma MicroRNA profiling reveals loss of endothelial MiR-126 and other MicroRNAs in type 2 diabetes. Circ. Res. 107. 10.1161/CIRCRESAHA.110.226357.

Zeng, Y., Cui, Z., Liu, J., Chen, J., and Tang, S. (2020). MicroRNA-29b-3p Promotes Human Retinal Microvascular Endothelial Cell Apoptosis via Blocking SIRT1 in Diabetic Retinopathy. Front. Physiol. 10. 10.3389/fphys.2019.01621.

Zhou, L., Li, F.F., and Wang, S.M. (2021). Circ-ITCH restrains the expression of MMP-2, MMP-9 and TNF-α in diabetic retinopathy by inhibiting miR-22. Exp. Mol. Pathol. 118. 10.1016/j.yexmp.2020.104594.

Zhu, Y., Yang, X., Zhou, J., Chen, L., Zuo, P., Chen, L., Jiang, L., Li, T., Wang, D., Xu, Y., et al. (2022). MiR-340-5p Mediates Cardiomyocyte Oxidative Stress in Diabetes-Induced Cardiac Dysfunction by Targeting Mcl-1. Oxid. Med. Cell. Longev. 2022. 10.1155/2022/3182931.

