## Supplementary material for "Transcriptomic Profile of Lin^-^Sca1^+^c-kit (LSK) cells in db/db mice with long-standing diabetes": Supplemental Information.pdf

### Supplemental Figures:

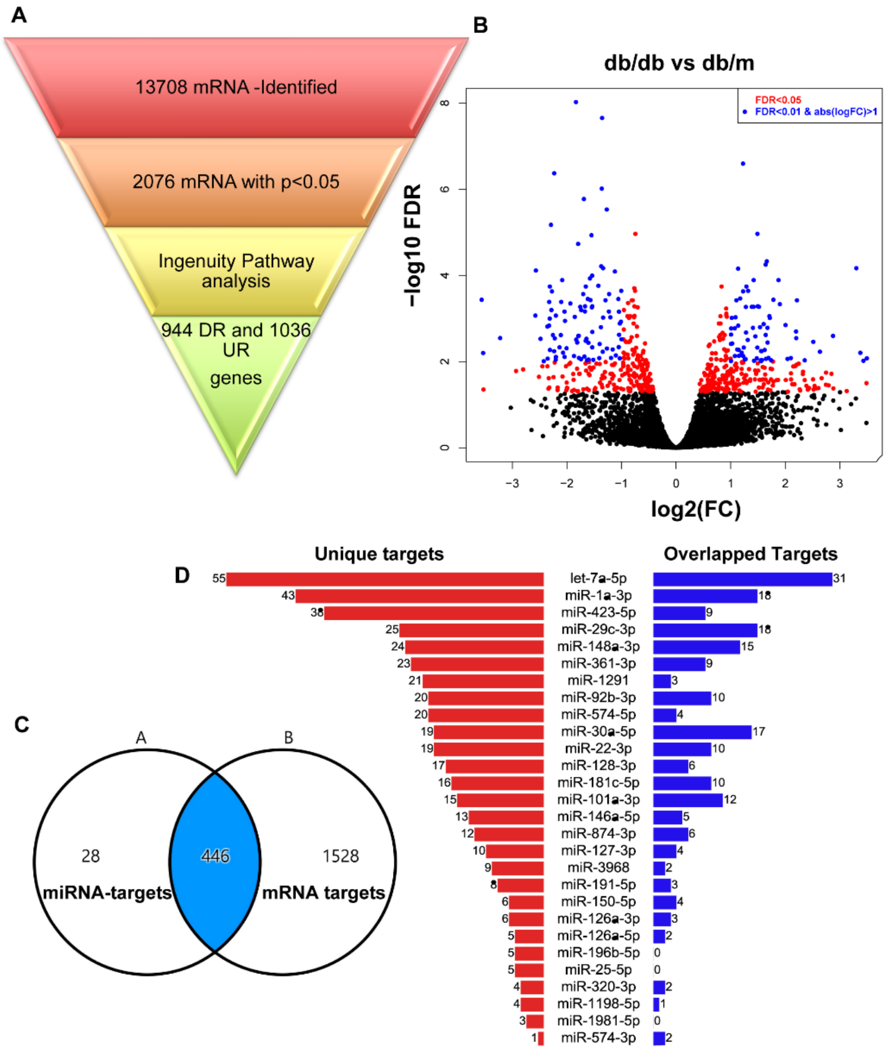


**Figure S1. Sequencing data analysis of HSCs in db/db mice.** The raw sequencing data for mRNA was subjected to various analyses. We found a total of 13708 mRNAs, out of which 2076 were statistically significant. **(A)** The schematic presentation of the steps followed for the data analysis for mRNAs. **(B)** The volcano plot represents all the differentially expressed mRNAs in db/db mice; blue and red dots were the significantly changed mRNAs. The top 35 (significant or p<0.05) miRNAs were subjected to MicroRNA filter analysis on IPA software and were overlayed with the mRNA targets found in our sequencing data. **(C)** Venn diagram depicting 446 common mRNA targets between miRNA-targets and mRNA targets obtained from the sequencing data. **(D)** MicroRNA target filter also showed that individual miRNA has some unique mRNA targets, and some mRNA targets were common among multiple miRNAs.

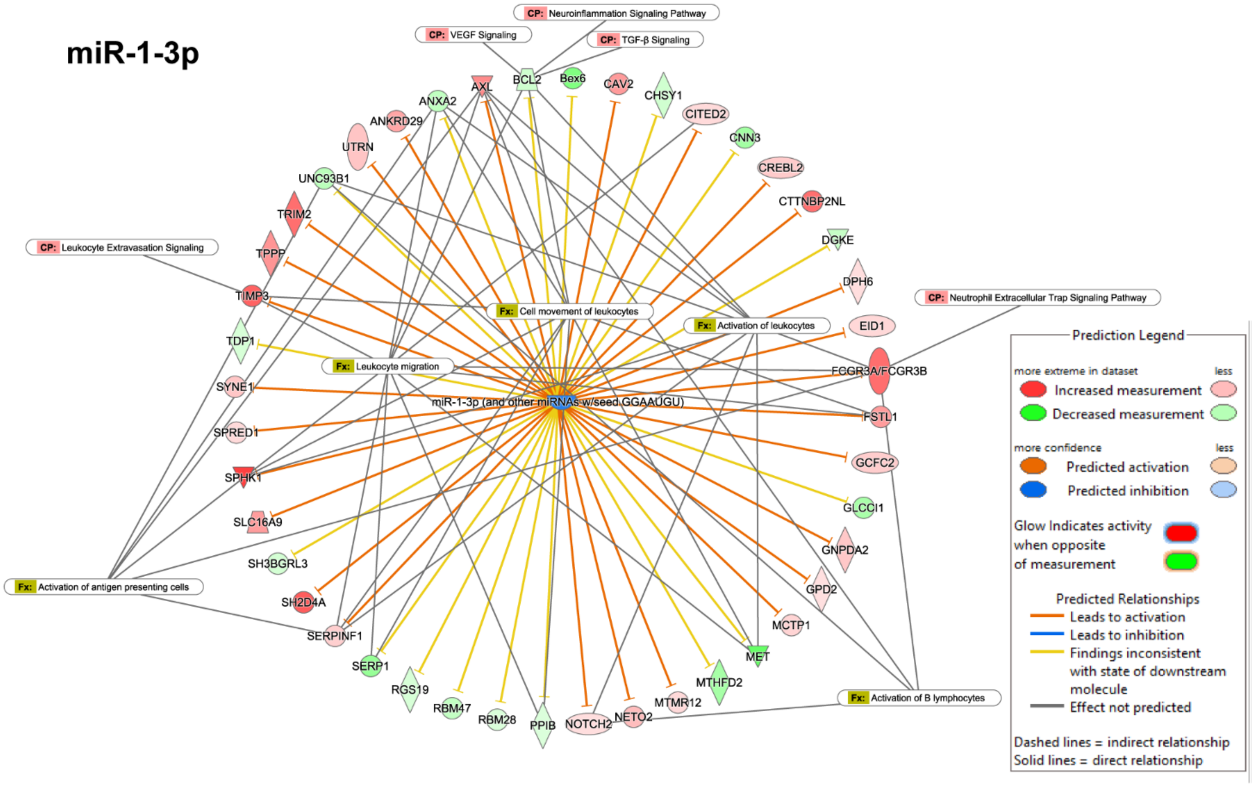


**Figure S2. IPA network analysis revealed possible involvement of neuroinflammatory and VEGF signaling downstream of miR-1-3p.** miR-1-3p was one of the top upregulated miRNAs observed in the HSCs of db/db mice. When subjected to IPA analysis, it was linked to leukocyte activation, migration, neuroinflammation, and VEGF signaling.

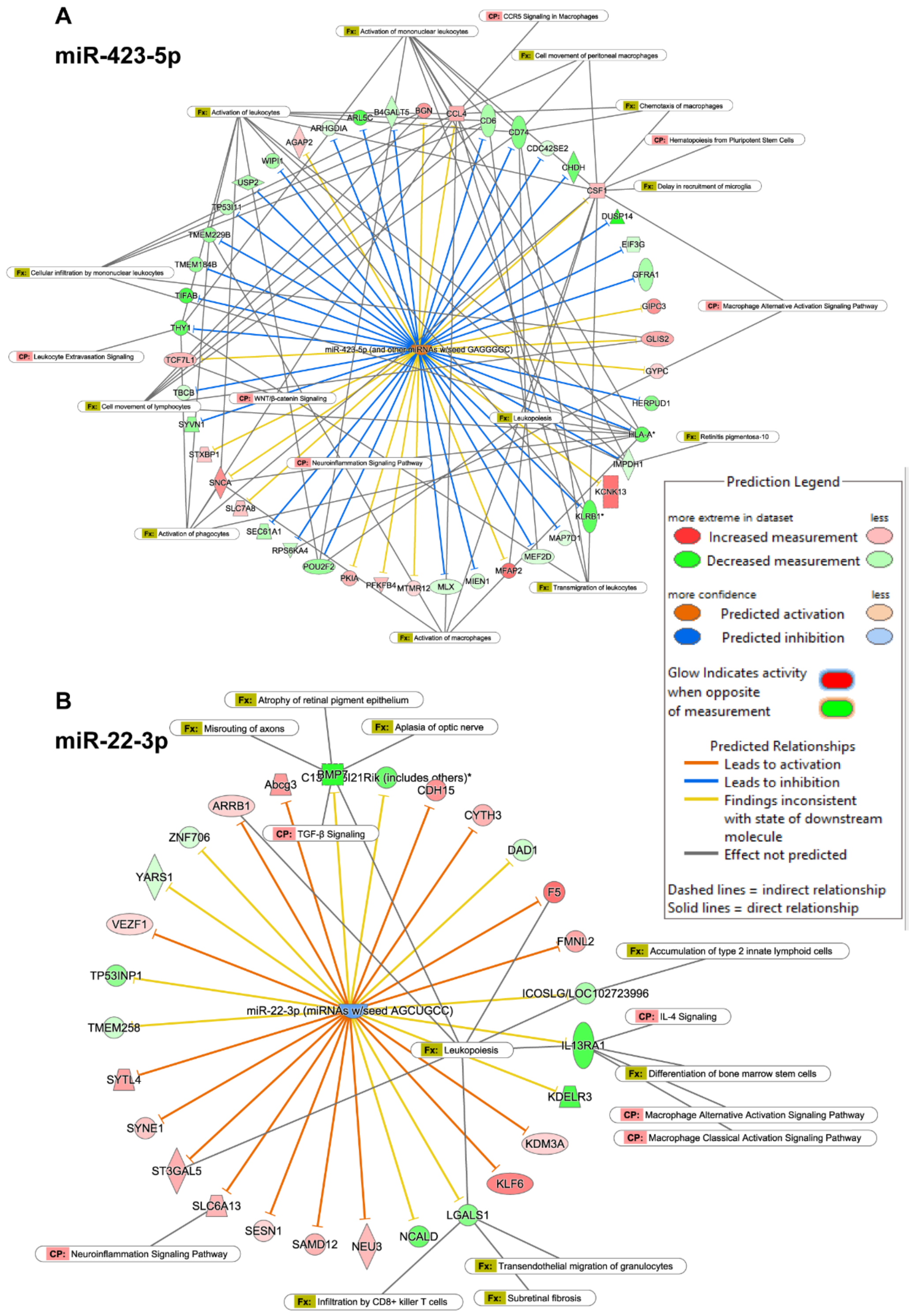


**Figure S3. IPA network analysis illustrating the top molecular pathways downstream of miR-423-5p and miR-22-3p.** miR-423-5p **(A)** and miR-22-3p **(B)** were among the top downregulated miRNAs, which showed the activation of macrophage signaling, neuroinflammation, microglia activation, and migration signaling.

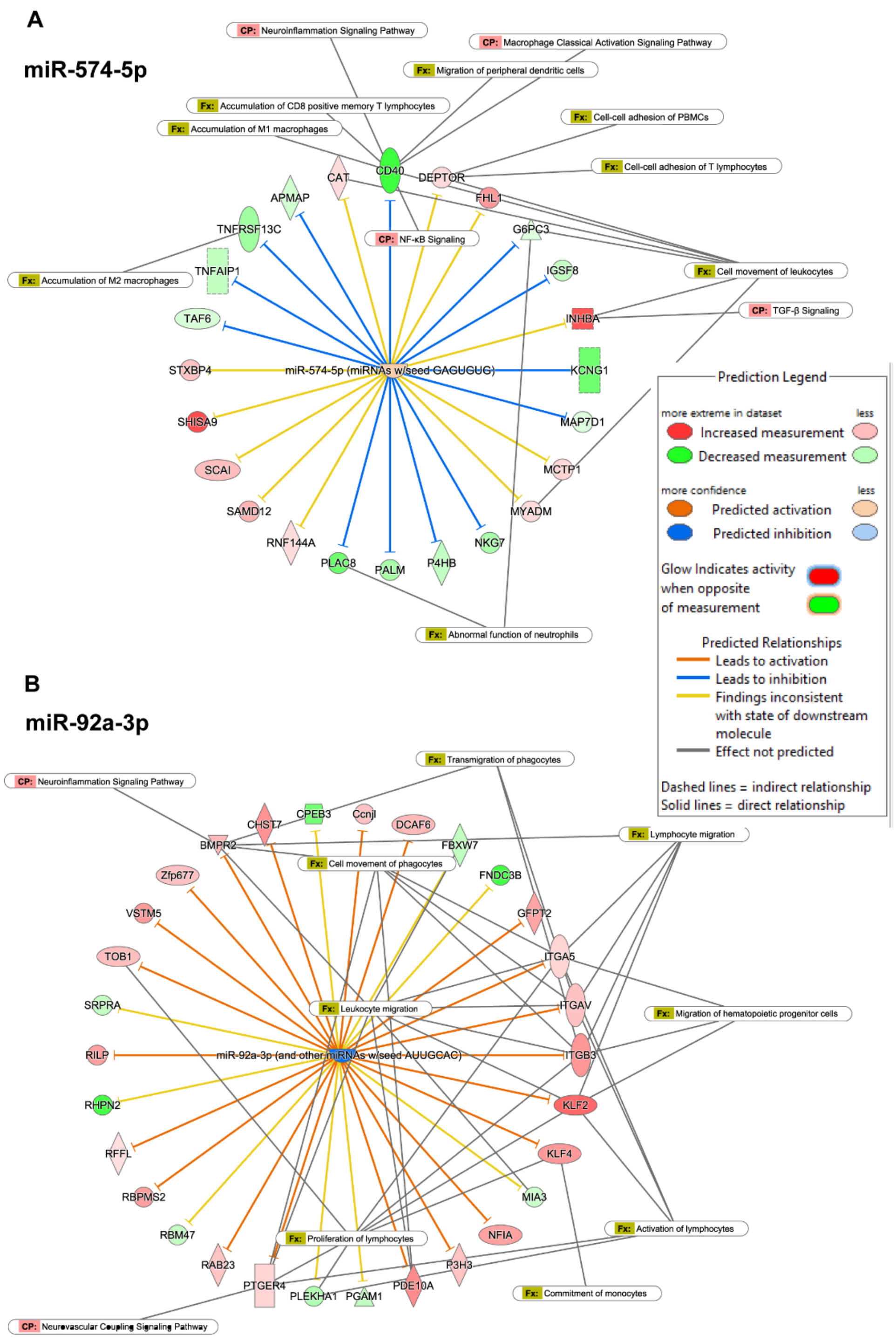


**Figure S4. Chronic diabetes activates inflammatory pathways at the transcriptional level in HSCs.** miR-574-5p and miR-92a-3p were also downregulated in the HSCs of db/db mice. **(A)**The network analysis for miR-574-5p revealed an upregulation of neuroinflammation, macrophage accumulation, and TGF-β signaling. **(B)** The downregulation of miR-92a-3p activated neuroinflammatory and leukocyte migratory pathways via upregulating integrins (*Itga5, Itgav, Itgb3*) and Kruppel-like factor 2 and 4 (*Klf2 and Klf4*).

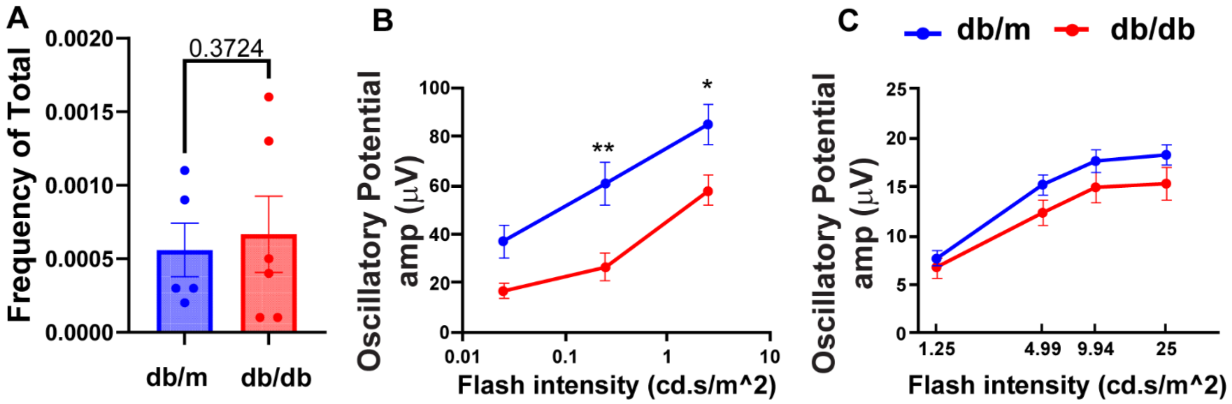


**Figure S5. Effect of chronic diabetes on retinal macrophage population and oscillatory potential.** **(A)**The macrophage (monocytes and neutrophils; CD11b$^+$CD45$^+$LyC$^+$LyG$^+$) population was higher in the db/db mice. The data were presented as mean ± SEM; N=6 and analyzed using a student's t-test.

**(B)**The average oscillatory potential showed a significant decline in the db/db mice at higher frequencies under scotopic conditions. **(C)** The average oscillatory potential was not significantly changed during photopic ERG. Data are expressed as mean ± SEM. **p<0.01; *p<0.01 show changes vs db/m mice. N=10 and the data were analyzed using two-way ANOVA followed by Sidak's multiple comparisons test.

**Table S1.** Top Analysis Ready MicroRNAs (10 Upregulated and 10 Downregulated)

| Sr No. | MiRNAs | Expression value | Expression Level |
|---|---|---|---|
| **1** | miR-1-3p (and other miRNAs w/seed GGAAUGU) | 1.841 | Upregulated |
| **2** | miR-126a-5p (and other miRNAs w/seed AUUAUUA) | 1.520 | Upregulated |
| **3** | miR-874-3p (miRNAs w/seed UGCCCUG) | 1.351 | Upregulated |
| **4** | miR-126a-3p (and other miRNAs w/seed CGUACCG) | 1.311 | Upregulated |
| **5** | miR-30c-5p (and other miRNAs w/seed GUAAACA | 1.277 | Upregulated |
| **6** | miR-101-3p (and other miRNAs w/seed ACAGUAC) | 1.022 | Upregulated |
| **7** | miR-181a-5p (and other miRNAs w/seed ACAUUCA) | 0.911 | Upregulated |
| **8** | miR-340-5p (miRNAs w/seed UAUAAAG) | 0.904 | Upregulated |
| **9** | miR-29b-3p (and other miRNAs w/seed AGCACCA | 0.783 | Upregulated |
| **10** | miR-196a-5p (and other miRNAs w/seed AGGUAGU) | 0.672 | Upregulated |
| **11** | miR-3968 (miRNAs w/seed GAAUCCC) | 2.011 | Downregulated |
| **12** | miR-22-3p (miRNAs w/seed AGCUGCC) | 1.510 | Downregulated |
| **13** | miR-574-5p (miRNAs w/seed GAGUGUG) | 1.503 | Downregulated |
| **14** | miR-148a-3p (and other miRNAs w/seed CAGUGCA) | 1.495 | Downregulated |
| **15** | miR-423-5p (and other miRNAs w/seed GAGGGGC) | 1.487 | Downregulated |
| **16** | miR-127-3p (miRNAs w/seed CGGAUCC) | 1.291 | Downregulated |
| **17** | miR-1971 (and other miRNAs w/seed UAAAGGC) | 1.245 | Downregulated |
| **18** | miR-361-3p (miRNAs w/seed CCCCCAG) | 1.209 | Downregulated |
| **19** | miR-320b (and other miRNAs w/seed AAAGCUG) | 1.120 | Downregulated |
| **20** | miR-1306-5p (miRNAs w/seed ACCACCU) | 1.100 | Downregulated |
